# Emerging Evidence for Poxvirus-Mediated Unfolded Protein Response: Lumpy Skin Disease Virus Maintains Self-replication by Activating PERK and IRE1

**DOI:** 10.1101/2022.06.27.497878

**Authors:** Jinlong Tan, Yinju Liu, Fan Yang, Guohua Chen, Yongxiang Fang, Xiaobing He, Zhongzi Lou, Huaijie Jia, Zhizhong Jing, Weike Li

## Abstract

The cytoplasmic replication of poxviruses requires extensive protein synthesis, challenging the capacity of the endoplasmic reticulum (ER). However, the role of the ER in the life cycle of poxviruses is unclear. In this study, we demonstrate that infection with the lumpy skin disease virus (LSDV), a poxvirus, causes ER stress *in vivo* and *in vitro*, further facilitating the activation of the unfolded protein response (UPR). Although UPR activation aids in the restoration of the cellular environment, its significance in the LSDV life cycle remains unclear. Furthermore, the role of ER imbalance for viral replication is also unknown. We show that LSDV replication is hampered by an unbalanced ER environment. In addition, we verify that the LSDV replication depends on the activation of PERK-eIF2α and IRE1-XBP1 signaling cascades rather than ATF6, implying that global translation and XBP1 cleavage are deleterious to LSDV replication. Our findings suggest that LSDV engages all UPR signaling sensors, and that activation of PERK and IRE1 sensors is indispensable to maintaining its own replication.

**IMPORTANCE:** Although numerous viruses cause ER stress and employ endogenous UPR components to control viral growth, there is no such evidence for poxviruses. Recent real-world epidemics of poxviruses such as monkeypox and LSDV indicated a lack of available coping strategies. Our findings show that LSDV encoding up to 156 open reading frames (ORFs) causes pressure to the stabilization of ER, triggers ER stress, and further promotes the activation of all three UPR signaling pathways. However, inhibiting PERK-eIF2α and IRE1-XBP1 was not conducive for LSDV replication. Since LSDV efficiently utilizes UPR components to assist its own replication, signal-blocking agents of PERK and IRE1 may be useful in the treatment of LSDV. More evidence for the efficacy of such therapy for LSDV, even monkeypox, could come from a clearer characterization of the ER stress-mediated viral replication process.

## INTRODUCTION

Eukaryotes are under long-term threat from viral invasion. Both DNA and RNA viruses rely on the steady operation of host organelles to provide the energy and substrates they need to sustain their replication (1, 2). Protein synthesis, modification, folding, and transport play critical roles in the viral life cycle. These functions are particularly linked to homeostasis of the endoplasmic reticulum (ER), the key site of protein synthesis (3–5).

When the ER is subjected to excessive protein demands from viruses, ER stress, an evolved conservative defensive mechanism, is triggered to restore the intracellular equilibrium (6, 7). The activation of three important signal sensors—protein kinase RNA-like ER kinase (PERK), inositol-requiring enzyme 1 (IRE1), and activating transcription factor 6 (ATF6)—initiates the unfolded protein response (UPR) (8–10) and the defensive mechanism (ATF6). These three UPR signal sensors are bound to molecular chaperones represented by GRP78 (also known as Bip) in a stable cellular environment, with little activation. Under stress, however, PERK, IRE1, and ATF6 are effectively activated when they dissociate from GRP78 (11, 12). The distinct intervention modes of the three signaling pathways characterize the potential of the UPR. Phosphorylation of PERK activates itself, causing phosphorylation of eIF2α to inhibit total protein translation. Meanwhile, ATF4 is activated to enhance CHOP production that leads to ER stress-mediated apoptosis (13, 14). IRE1 is also triggered by autophosphorylation that stimulates XBP1 cleavage and enhances the transcription of molecular chaperones (15, 16). ATF6 is activated by cleavage from full-length 90-kDa ATF6 (ATF6-FL) to activated N-terminal 50-kDa ATF6 (ATF6-N); this ameliorates overloaded protein transport to the Golgi apparatus (17, 18).

Surprisingly, distinct viruses appear to utilize unique UPR strategies to regulate their replication (3, 19–21). In other words, host cells may flexibly respond to viral invasion through the UPR. Similarly, replication processes of poxviruses, being DNA viruses with large genomes, have not been well defined in relation to ER stress.

In the present study, the relationship between lumpy skin disease virus (LSDV, a member of the poxvirus family) and ER stress was investigated. LSDV belongs to the genus *Capripoxvirus* characterized by large genomes of more than 150 kbp and 156 open reading frames (ORFs) (22). The virus is a member of the *Chordopoxvirinae* subfamily of the *Poxviridae* family. In recent years, some studies (23–25) have claimed that membrane components of poxviruses originated from the ER, implying that the ER most likely assembles recruited viral proteins. These extra protein processing requirements likely overburden the ER, resulting in ER stress. Analysis of the ER disturbance may provide insights into viral membrane dynamics and ER homeostasis during poxvirus infection through biochemical investigations of the membrane origin and ER function in the poxvirus life cycle.

Our results demonstrate a breakthrough in ER stabilization during LSDV infection. Furthermore, UPR-related chemicals may contribute to the treatment of LSD.

## RESULTS

### LSDV infection induces a characteristic phenotype *in vivo*

To establish effective LSDV infection *in vivo*, the supernatant containing LSDV was intradermally multipoint injected into three cows. Meanwhile, three other cows were similarly injected with DMEM. At 14 days post-infection (dpi) with LSDV, we observed typical lumpy skin growing on the body surfaces of the Holstein cows. However, cows managed with DMEM displayed no variation (Fig. 1A). In addition, multiple rectal temperature tests showed that LSDV infection caused varying degrees of temperature rise in the cows that could be transient or long-term, depending on individual differences. However, the temperature of control cows did not change significantly (Fig. 1B). During infection, LSDV was monitored multiple times using an LSDV ELISA kit produced by our institute. We detected LSDV-positive blood samples in LSDV-infected cows at 14 dpi, while no cows tested LSDV positive in the early stages of infection, suggesting that early latent infection of LSDV might not be conducive to surveillance. All control cows presented no LSDV positives during the trial (Fig. 1C). H&E staining was performed to investigate the effects of LSDV infection on skin tissues. Histological observation revealed that nodular skin tissues were present in a large area of the dermis showing cell necrosis, shallow dermis visible collagen fiber necrosis (black arrow in the figure) dotted with inflammatory cell infiltrate (red arrows), deep dermis collagen fiber necrosis, and pieces of dead cells (yellow arrow) accompanied by a small amount of inflammatory cell infiltration (blue arrow). Conversely, none of these pathological changes were found in the skin of uninfected cows or in normal skin adjacent to the nodules (Fig. 1D). Collectively, these results demonstrated the successful construction of an LSDV infection model.

**Figure 1.**
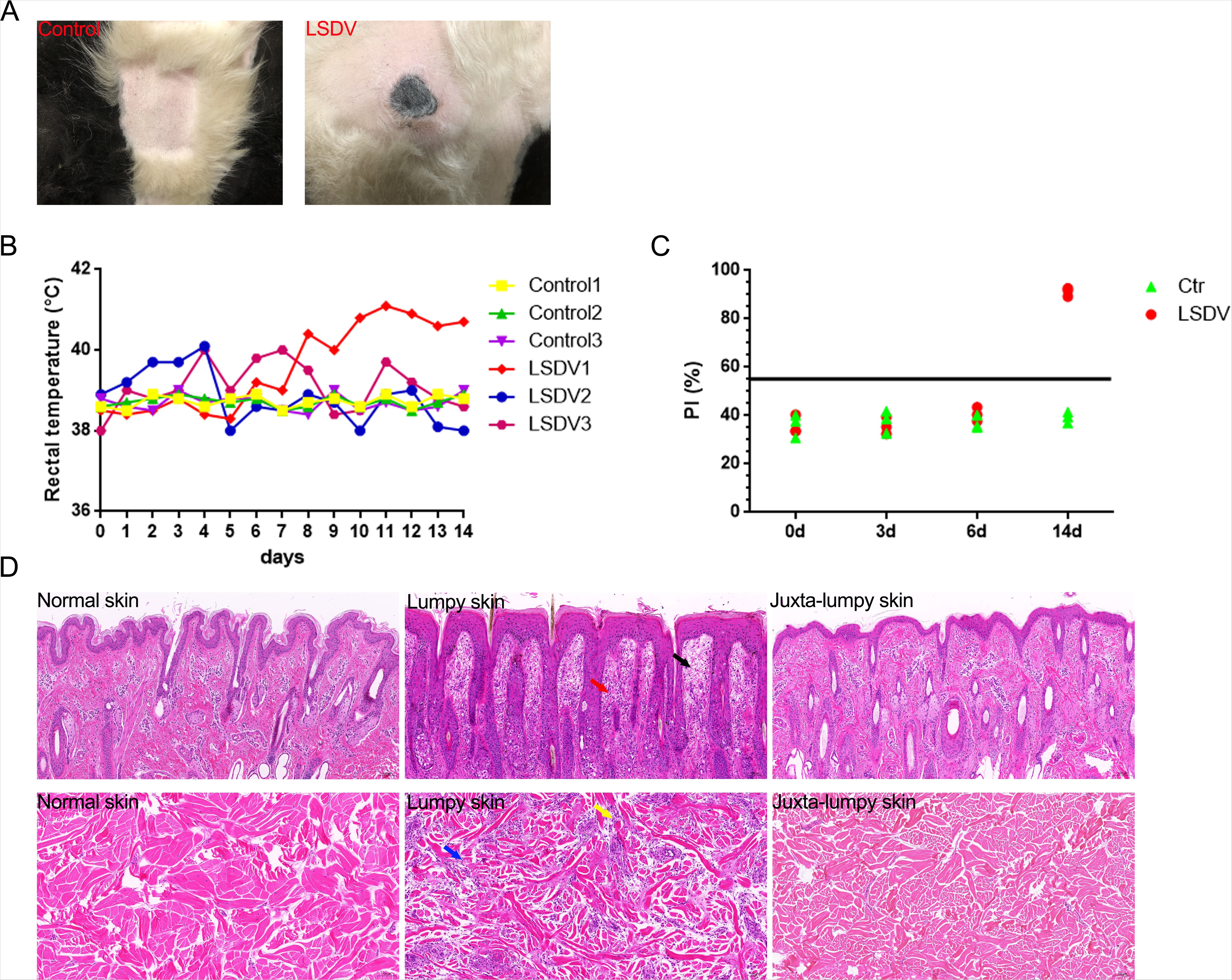
LSDV infection elicits a characteristic phenotype *in vivo*. (A) Images of normal skin from control cows and lumpy skin from LSDV-infected cows. (B) The rectal temperature monitoring of cows after control management and LSDV intervention. (C) Serological identification of LSDV infection via ELISA, n = 3. (D) The histopathological changes (H and E staining) in the two groups of cows were observed.

### LSDV infection elicits ER ultrastructural abnormalities and ER stress *in vivo*

To explore whether LSDV infection was associated with an unbalanced ER, transmission electron microscopy (TEM) was conducted to examine structural variation in the ER of infected or uninfected skin fibroblasts. Mature and immature virions were observed in LSDV-infected skin fibroblasts (yellow arrow in Fig. 2A), demonstrating that cells were undergoing LSDV infection. Compared with the normal skin fibroblasts, LSDV-infected skin fibroblasts revealed increased ER and a dilated ER lumen (orange arrow in Fig. 2A), indicating that LSDV infection may disturb ER progression and further trigger ER stress. Notably, mitochondrial swelling was also observed in LSDV-infected skin fibroblasts (green arrow in Fig. 2A), suggesting that LSDV infection mediated mitochondrial damage. to further investigate the association of LSDV infection with ER stress, a primary marker GRP78, also known Bip, was used to characterize ER homeostasis. Compared with normal skins of control cows, lumpy skins revealed a high expression of GRP78 (Fig. 2B). Additionally, fluorescence co-staining of GRP78 and ORF122 showed that LSDV and GPR78 were abundantly distributed in the lumpy skin, especially in the dermis, as characterized by fluorescence results of longitudinal cutting and crosscutting, respectively. However, there was no significant positive distribution of GRP78 in the juxta-lumpy skin or in the normal skin of control cows, and only a small amount of LSDV infection was found in the juxta-lumpy skin (Fig. 2C). Furthermore, IHC verified the results as described by fluorescence co-staining (Fig. 2D). In a word, LSDV infection triggered ER stress *in vivo*.

**Figure 2.**
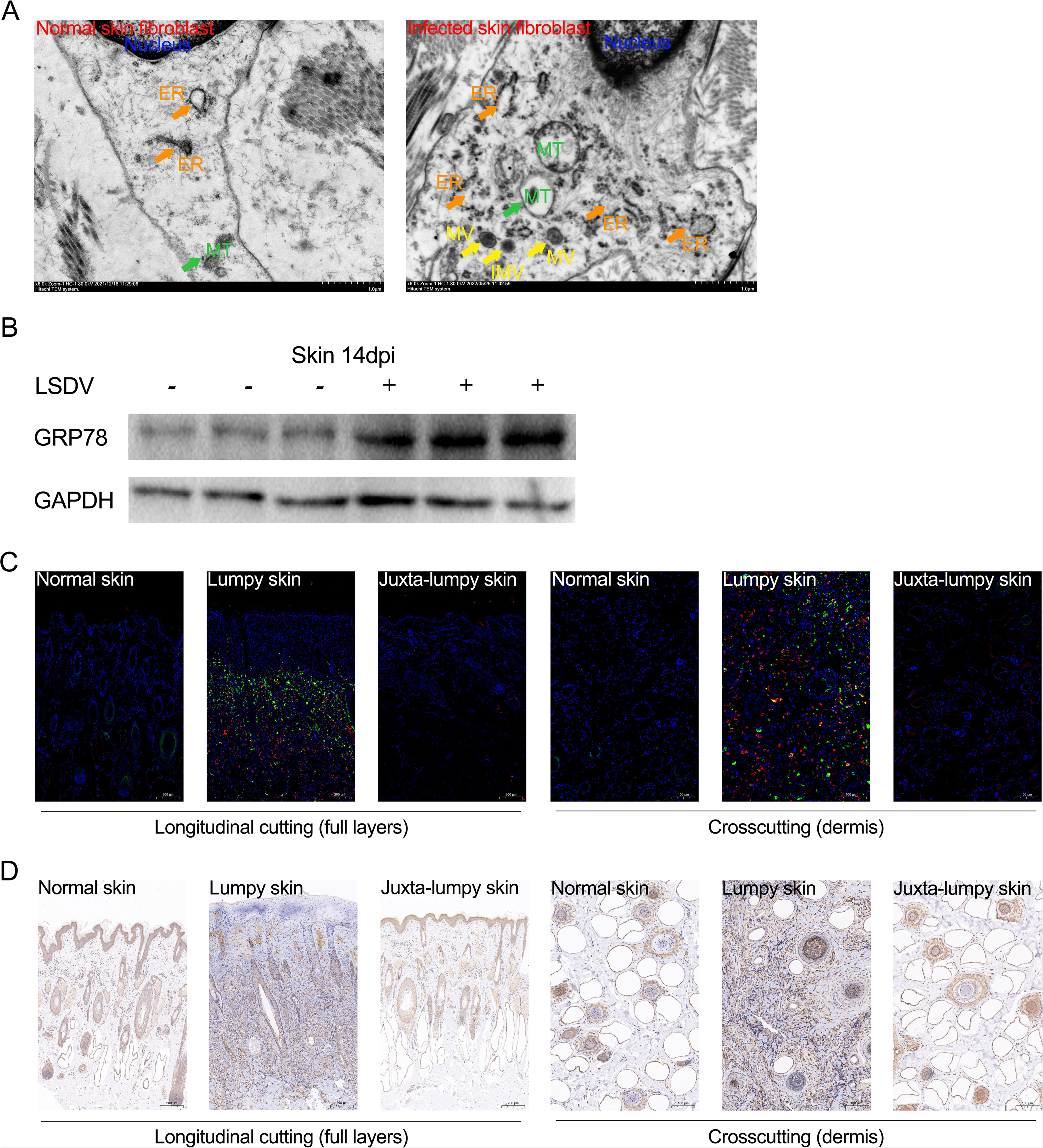
Identification *in vivo* of LSDV infection inducing ER stress. (A) Ultrastructural observations of uninfected and LSDV-infected skin fibroblasts by TEM. (B) Skin sample immunoblot assay of GRP78 in control group and LSDV-infected group cows, n = 3. (C) The normal skin of control cows, lumpy skin of infected cows, and normal skin juxta-lumpy skin are labeled (full layers and dermis; Blue denotes DAPI, green indicates GRP78, and red shows ORF122) by IFA. (D) IHC again identified a positive distribution of GRP78.

### LSDV infection activates all three UPR signaling pathways *in vivo*

Generally, ER stress is accompanied by differential activation of UPR signaling pathways. In this experiment we discovered that all three UPR signals were activated in the infected lumpy skin compared to the skin of control cows. In detail, PERK phosphorylation promoted the activation of PERK signals that further facilitated phosphorylation of eIF2α and initiated translation attenuation. Similarly, IRE1 was phosphorylated for activation and intensified the expression of the downstream cleaved form XBP1 (sXBP1), contributing to the transcription of molecular chaperones and recovering ER stabilization. When activated, ATF6 is translocated to the Golgi apparatus to achieve the cleavage of the ATF6 N-terminal form (ATF6-N), thus participating in the nuclear regulation and restoration of ER homeostasis. We observed that LSDV infection triggered the expression of activated ATF6-N, indicating that an attempt to restore the imbalance of ER by activating ATF6 during LSDV infection (Fig. 3). Overall, the LSDV infection activated all three UPR signals. In other words, these three strategies may contribute to the recovery of LSDV-induced ER stress.

**Figure 3.**
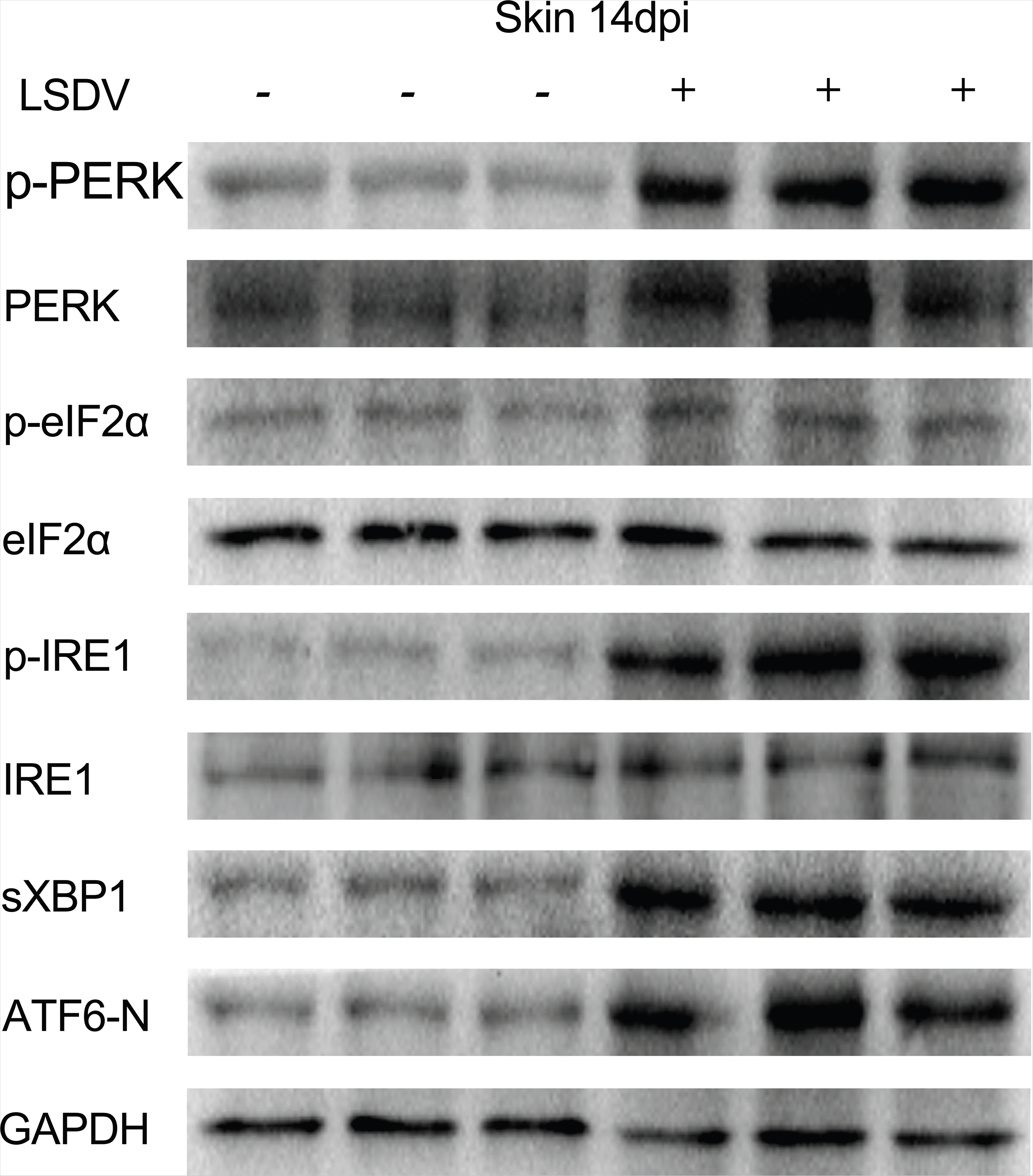
Skin immunoblot assay of UPR signal in the control group and LSDV-infected group cows. n=3.

### LSDV infection triggers ER stress *in vitro*

To validate that LSDV infection induced ER stress *in vitro*, the transcriptional and translational levels of GPR78 were monitored in the LSDV-infected or uninfected BEF or MDBK cells (cultured in 6-well plates). The results suggested that neither BEF nor MDBK cells revealed variation in GRP78 transcriptional levels at 12 hours post infection (hpi) and 24 hpi, while both cells showed a significant transcriptional increase at 48 hpi and 96 hpi (Fig. 4A and B). Similarly, we observed no change in the expression of GRP78 protein in either cells at 12 hpi and 24 hpi, regardless of whether cells were infected with LSDV (Fig. 4C and D). All of these results demonstrated that LSDV infection triggered ER stress *in vitro*.

**Figure 4.**
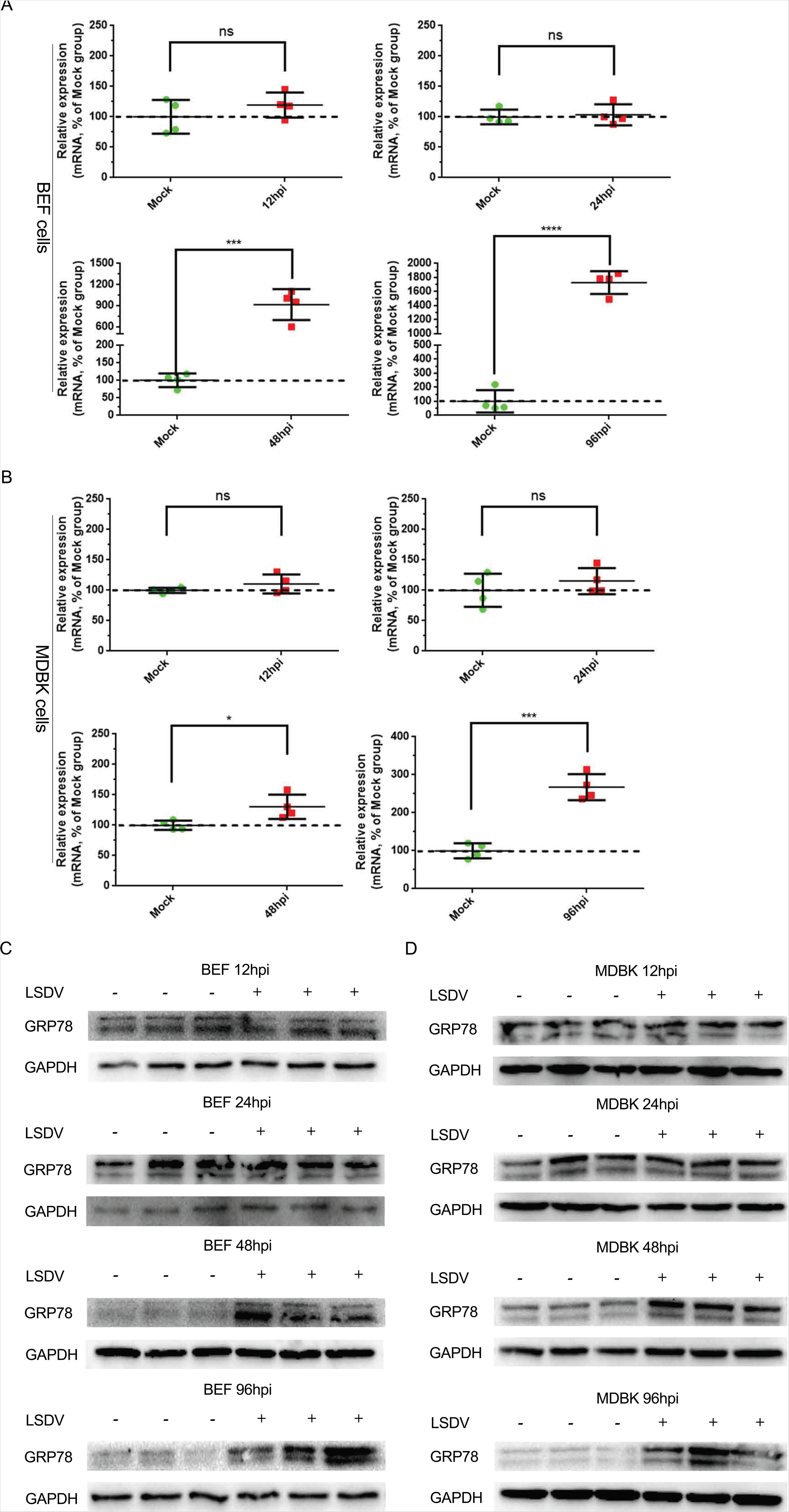
(A and B) Detection of GRP78 relative mRNA levels in mock infected or infected MDBK and BEF cells with LSDV (MOI=1) for 12 h, 24 h, 48 h, and 96 h (n = 4). The dotted lines represented the mock control, and the same below. (C and D) GRP78 immunoblot assay in mock infected or infected MDBK and BEF cells with LSDV (MOI = 1) for 12 h, 24 h, 48 h, and 96 h (n = 3).

### LSDV infection activates all three UPR signaling sensors *in vitro*

Next, we examined whether the UPR strategies were applied *in vitro*. The protein levels of UPR signaling pathways were monitored in both BEF and MDBK cells (cultured in 6-well plates) at different time points. Immunoblotting results showed that LSDV did not promote the activation of PERK, IRE1, or ATF6 in both types of cells at 12 hpi and 24 hpi. Conversely, all three UPR signals were activated in LSDV-infected cells at 48 hpi and 96 hpi, further facilitating the phosphorylation of eIF2α and cleavage of XBP1 and N-terminal ATF6 (Fig. 5A and B). These results indicated that LSDV infection triggered the activation of UPR signaling sensors *in vitro*.

**Figure 5.**
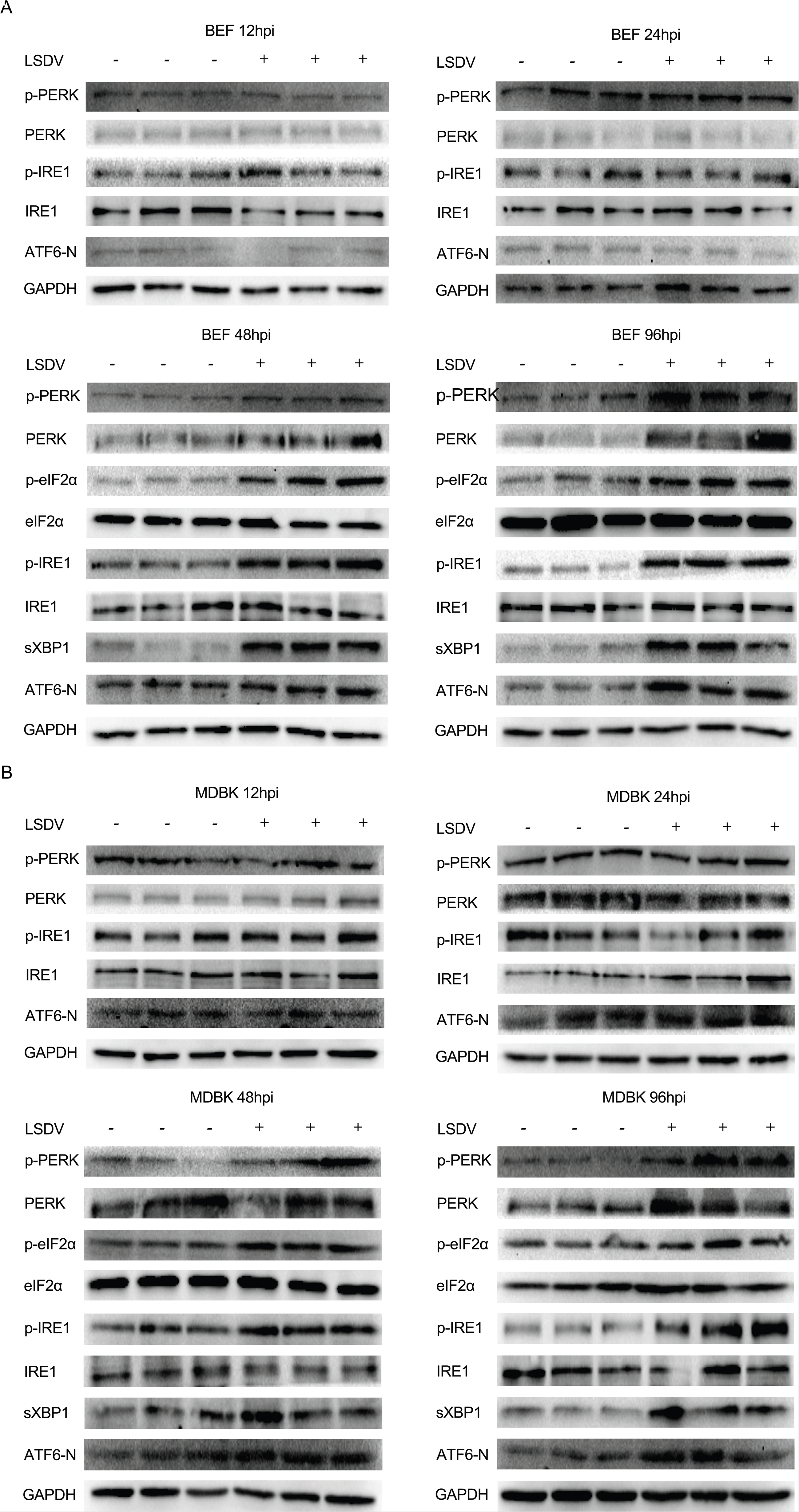
(A and B) Immunoblot assay of UPR signaling pathways in mock infected or infected MDBK and BEF cells with LSDV (MOI = 1) for 12 h, 24 h, 48 h, and 96 h (n = 3).

### Tunicamycin (TM)-induced ER stress is detrimental to LSDV replication in BEF and MDBK cells

Tunicamycin is an effective ER stress inducer causing accumulation of unfolded proteins in the ER lumen. Here, TM was utilized for investigating the role of ER stress in LSDV replication. After treatment with different TM concentrations (0.01, 0.05, and 0.25 μg/mL) for 24 h, CCK-8 was detected using 96-well plates to uncover an appropriate concentration (0.01 μg/mL) that did not affect cell activity in either type of cells (Fig. 6A). Subsequently, 0.01 μg/mL TM and an equal amount of DMSO were suppled to both cells for 24 h treatment, causing the ER stress environment and mock management, respectively (cells were spread onto 6-well and 12-well plates, respectively). Next, cells treated with TM and DMSO were infected with LSDV (MOI = 1) for 48 h. After that, WB, IFA, absolute quantification PCR, and virus titration were performed to explore the effect ER stress on LSDV replication. The results showed that TM management increased the protein expression of the ER stress marker GRP78 in both cells, while suppressing ORF122 expression (Fig. 6B). Prior to that, a band of ORF122 around 20 kDa was specifically detected in LSDV-infected cells (see Fig. S1 in the supplemental material). To further determine the role of LSDV infection in viral distribution, IFA was conducted to label ORF122 expression. Before IFA labeling, the specificity of ORF122 was identified, and no fluorescence signal of ORF122 was detected in negative BEF or MDBK cells (Fig. S2). We observed that TM treatment suppressed cellular lumpy CPE and decreased ORF122 distribution in both cells compared with DMSO management (Fig. 6C). Consistent with these findings, TM treatment notably inhibited viral copy numbers and decreased virus titer in both cells compared with DMSO management. Altogether, the induced ER stress was detrimental to LSDV replication *in vitro*.

**Figure 6.**
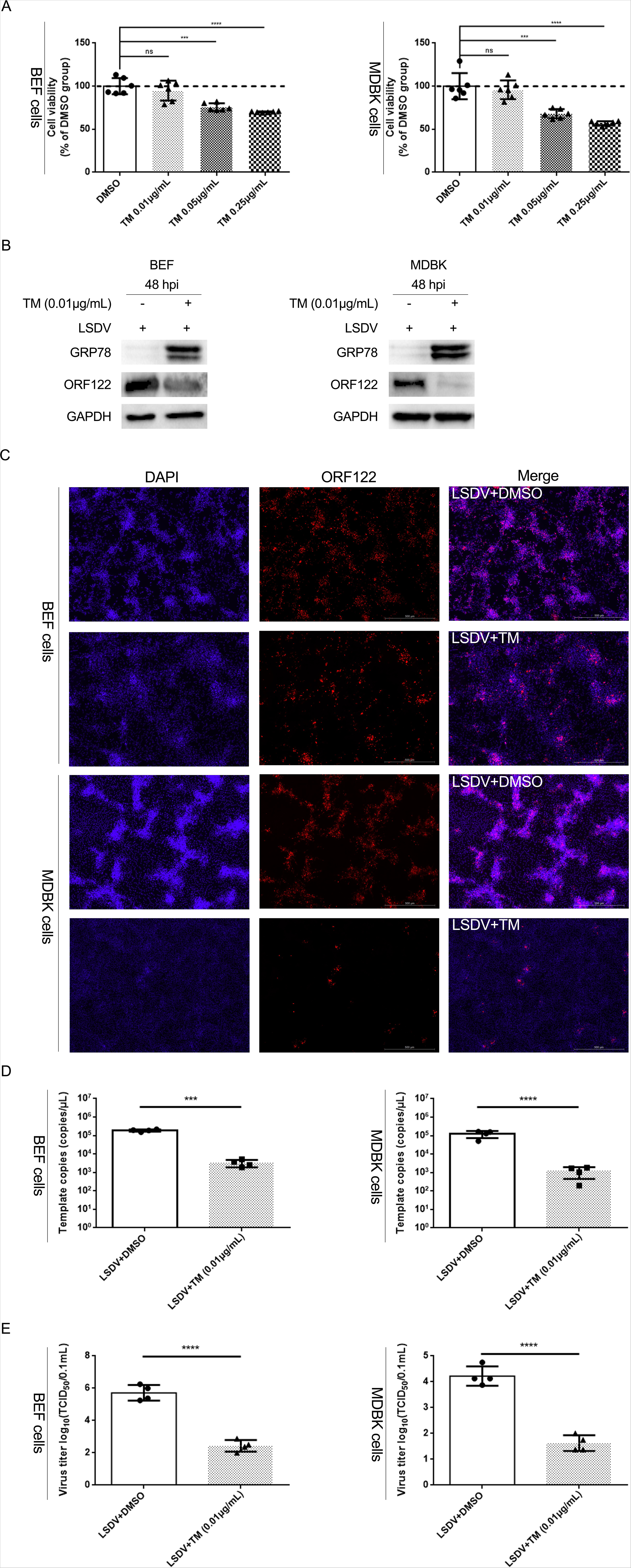
ER stress suppressed LSDV replication in both MDBK and BEF cells. Both cells were pretreated with DMSO and TM for 24 h and then infected with LSDV (MOI = 1) for 48 h. (A) The effect of TM on cell viability in both types of cells, n = 6. (B) Western blotting was performed to detect GRP78 and ORF122 expression. (C) Cellular IFA revealed the effect of TM on ORF122 distribution and lumpy CPE. (D) Viral copy number was detected by absolute quantification, n = 4. (E) Collected viral supernatant in both cells was titrated in LT cells, n = 4.

### RNA interference suggests that activation of PERK and IRE1 contributed to LSDV replication

Cells suppressed translation initiation by PERK, promoted ER chaperones transcription by IRE1-mediated cleavage of XBP1 into the nucleus, and regulated cascade signaling by N-terminal ATF6 cleavage of the Golgi apparatus, thus alleviating the pressure on the ER caused by unfolded protein overload. There is no doubt that activation of all three UPR signaling sensors facilitates recovery of LSDV-induced ER stress. It was worth investigating whether the activation of all three signals contributed to LSDV replication. Here, three siRNAs for each UPR signal sensor were screened to uncover a representative siRNA in both cells (cultured in 6-well plates). As shown in Fig. 7A, si-PERK3, si-IRE1-1 and si-ATF6-2 were utilized in subsequent experiments (cells were grown in 12-well plates) due to their capacity to interfere with the activation of their respective signals. We found that the CPE caused by LSDV infection was alleviated and that the distribution of ORF122 was reduced after treatment with si-PERK and si-IRE1 in both cells compared with the si-Control treatment, while si-ATF6 treatment did not affect the CPE or distribution of ORF122 caused by LSDV infection in either type of cells (Fig. 7B). Similarly, RNA interference to PERK and IRE1 markedly suppressed the number of virus copies and the virus titer in both types of cells compared with the si-Control (Fig. 7C and D), while RNA interference to ATF6 revealed no potency.

**Figure 7.**
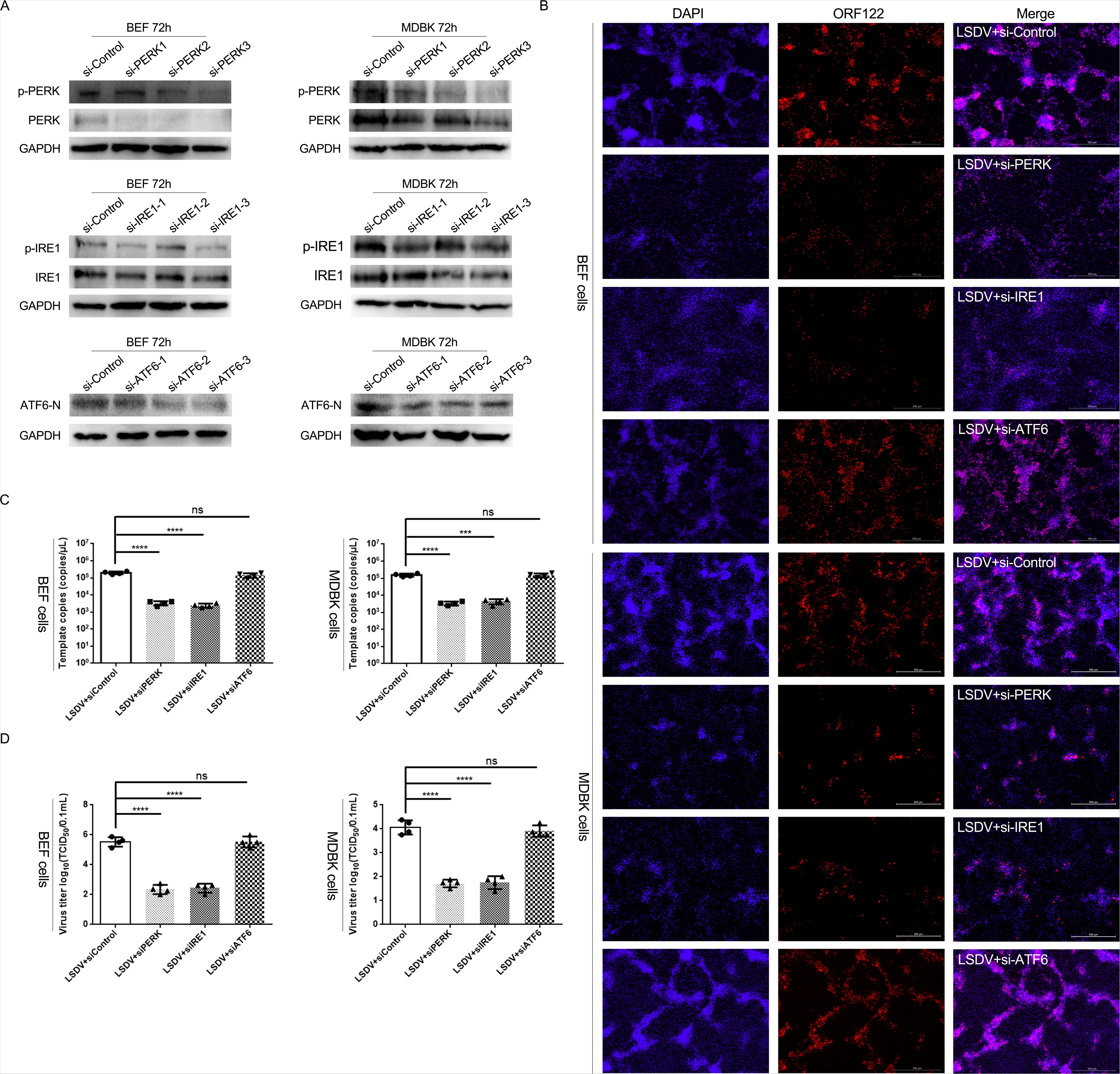
Three UPR signaling siRNAs demonstrated the activation of PERK and IRE1 pathways contributing to LSDV replication. (A) After treatment with 9 siRNAs for 72 h, the most effective UPR siRNAs that could block distinct signaling sensors were screened by western blotting. (B) Both MDBK and BEF cells were pretreated with control-siRNA and UPR-siRNAs for 24 h and then infected with LSDV (MOI=1) for 48 h. Western blots assessed the effect of the three most effective siRNAs on ORF122 expression. (C) Cellular IFA revealed the effect of three UPR siRNAs on ORF122 distribution and lumpy CPE, n = 4. (D) Viral copy number was evaluated after treatment with three siRNAs. (E) Cellular supernatant was titrated in LT cells, n=4.

### Chemical suppression of UPR demonstrates that LSDV replication relies on PERK-eIF2α and IRE1-XBP1 instead of the ATF6 pathway

To further confirm that suppression of the PERK and IRE1 pathways was detrimental to LSDV replication whereas ATF6 inhibition was not, three reagents (GSK2606414 [PERKi], 4μ8C, and Ceapin-A7) were used to respectively intercept PERK, IRE1, and ATF6 signaling activation. GSK2606414, a highly selective inhibitor of the PERK sensor, is commonly used to block eIF2α activation and further inhibit translation initiation. The reagent 4μ8C is a highly effective IRE1 blocker that can inhibit the cleavage of XBP1. Ceapin-A7 is also a potent inhibitor that can block the cleavage of N-terminal ATF6. After treating BEF and MDBK cells with different concentrations of these chemicals for 24 h, the cell viability was assessed with CCK-8 using 96-well plates. Suitable chemical concentrations (PERKi 10 μM, 4μ8C 50 μM and Ceapin-A7 10 μM) that did not affect cell viability were screened for subsequent experiments (Fig. 8A). After treatment of both cells with DMSO and the three separate reagents, cells cultured in the 6-well plates were infected with LSDV (MOI=1) for 48 h. We observed that PERKi suppressed the phosphorylation of PERK and further suppressed the phosphorylation of eIF2α, indicating that the chemical PERKi effectively blocked the translational suppression. Likewise, 4μ8C blocked the phosphorylation of IRE1 and further facilitated the cleavage suppression of XBP1. Ceapin-A7 also inhibited the cleavage of N-terminal ATF-6. Notably, PERKi and 4μ8C suppressed the expression of ORF122, while Ceapin-A7 did not (Fig. 8B). Consistent with the above results, the supplementing of PERKi and 4μ8C decreased the distribution of virus protein ORF122 and suppressed lumpy CPE in both types of cells, while adding Ceapin-A7 had no impact (Fig. 8C). Also, the addition of PERKi and 4μ8C significantly suppressed LSDV copy number and decreased virus titer compared with DMSO controls. Conversely, treating with Ceapin-A7 showed no effect on LSDV copy number or virus titer (Fig. 8D and E). Taken together, blocking PERK and IRE1 signaling pathways was not conducive to LSDV replication.

**Figure 8.**
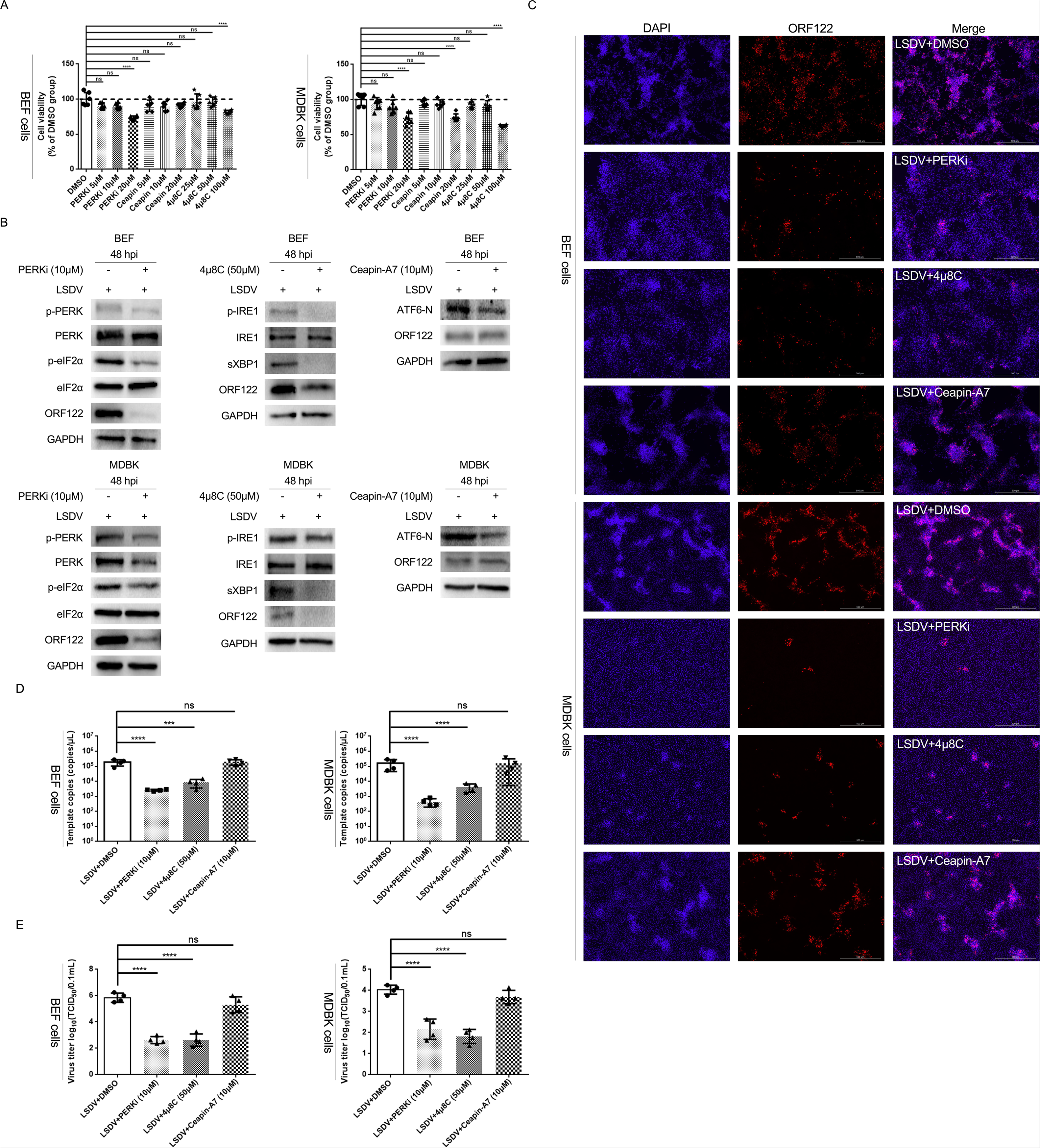
The intervention of UPR chemicals revealed that the activation of PERK and IRE1 was responsible for LSDV replication. (A) After treatment for 24 h, the effects of GSK2606414, 4μ8C, and Ceapin-A7 on cell viability were detected, n = 6. (B) Immunoblot assay of the suppression potency after intervention with inhibitors for 24 h. (C) Both MDBK and BEF cells were pretreated with DMSO and distinct chemicals for 24 h, then infected with LSDV (MOI = 1) for 48 h. Cellular IFA revealed the effects of the three reagents on ORF122 distribution and lumpy CPE. (D) Viral copy number discrepancy between DMSO-pretreated and inhibitor-pretreated cells, n = 4. (E) Collected supernatant was titrated in LT cells, n = 4.

## DISCUSSION

Numerous potential factors increase the risk of ER stress, including the large protein requirement of viruses, the excessively fast protein synthesis rate in the single replicating cycle, ER damage, and compensatory hypofunction of the ER due to damage of other organelles (26–28).

We demonstrated that when LSDV infection was present, ER stress was produced, and all three UPR branches were activated. Induction of ER stress and activation of the UPR may mediate various life stages in distinct viruses. ER stress was previously thought to play a role in viral replication. One study found that ER stress increased viral multiplication via ER stress-mediated autophagy (19). Likewise, another study found that the ER stress environment facilitated the replication of porcine circovirus type 2 (PCV2) (29). Other studies, however, suggested that ER stress was detrimental to viral replication. A study showed that chemically induced ER stress effectively inhibited coronavirus replication, implying that the creation of ER stress compounds might aid in the treatment of the growing SARS-CoV-2 pandemic (30). Similarly, a prior study found that therapy with an ER stress chemical inducer inhibited the replication of porcine hemagglutinating encephalomyelitis virus (PHEV) (31). These findings demonstrate that the ER stress affects the replication of various viruses in different ways. In our study we observed that TM-induced ER stress was detrimental to the replication of LSDV, implying that ER stress molecules might aid in the treatment of LSDV-mediated clinical symptoms. Clearly, enhancing ER stress is a candidate strategy for preventing LSDV infection, whereas a stronger ER stress could induce abnormal processes such as cellular death and autophagy (32, 33). As a result, further evaluation studies need to be conducted before LSD may be treated therapeutically with an ER stress inducer.

The induction of ER stress further facilitated UPR activation (34–36). It is particularly intriguing to observe how different UPR signaling pathways impact viral life progression. The replication of Kaposi’s sarcoma-associated herpesvirus (KSHV) depends on the activation of all three UPR sensors (3). Two investigations have shown that all three UPR signals are required for effective CSFV replication (37, 38). The activation of PERK and IRE1 increases the replication of porcine epidemic diarrhea virus (PEDV), but ATF6 has no effect on PEDV replication (39). Seneca Valley virus (SVV) induces autophagy via PERK and ATF6 pathways and further facilitates self-replication (40). Although the UPR is a common host response to protein accumulation, the data imply that it plays diverse roles in different viral life cycles. According to our results, we present an illustration (Fig. 9) of the detailed process of ER stress-mediated LSDV infection. After entering the cytoplasm, LSDV results in the accumulation of unfolded or misfolded proteins in the ER lumen. In order to restore ER homeostasis, Bip (GRP78) is translocated to the unfolded or misfolded protein for the signal transcription of folding, modulation, or trafficking. Following Bip’s dissociation from PERK, IRE1, and ATF6, PERK increases eIF2α phosphorylation and inhibits global translation, allowing LSDV replication and infection to proceed. In addition, IRE1 phosphorylation triggers the cleavage of downstream XBP1, facilitating LSDV replication and infection. Notably, ATF6 is also activated, although it appears to not enhance LSDV’s infectious potency. Overall, our results show that PERK and IRE1 activation aids LSDV infection.

**Figure 9.**
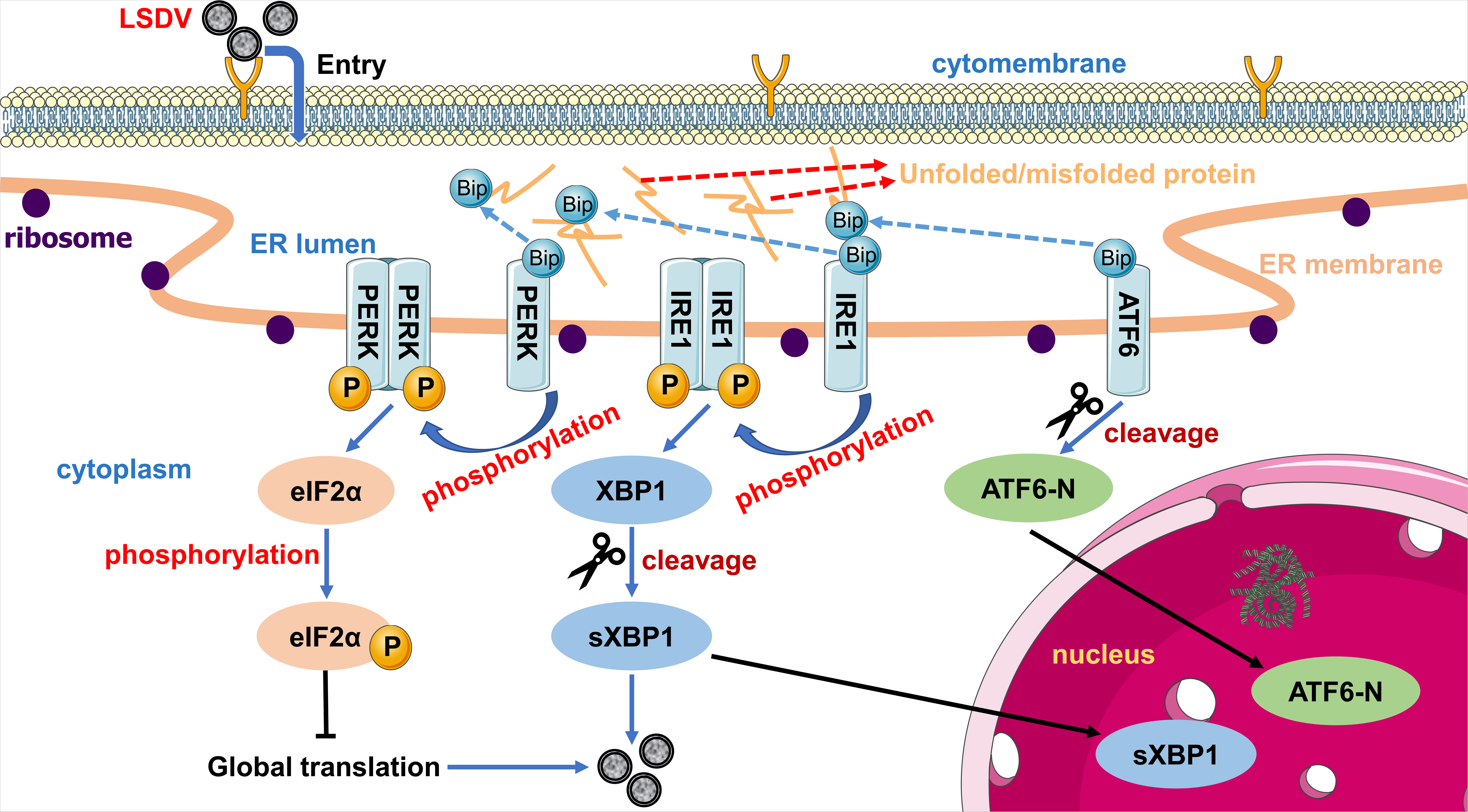
Illustration of ER stress-mediated LSDV infection.

Suppression with various compounds appears to be a potential UPR pharmacological therapy. It has been shown that pharmacological suppression of ATF6 and IRE1α branches can effectively decrease coronavirus infection (41), indicating that the UPR is a viable antiviral target. Another study found that a UPR chemical inhibitor (PERK) significantly improved clinical symptoms of Middle East respiratory syndrome coronavirus (MERS-CoV) (42), indicating the prospect of UPR chemicals being used to treat viral infections. In our study, we also demonstrated that UPR chemicals contributed to inhibiting LSDV infection *in vitro*. For assurance of UPR treatment of LSD, further *in vivo* evidence needs to be provided, including evaluation of clinical symptoms, disease prognosis, and adverse reactions.

Skin, a natural barrier between the outside world and the inside of the body, is composed of epidermis, dermis, and subcutaneous tissue. Skin may be able to successfully withstand external chemical, physical, and pathogenic stimuli (43–45). ER stress has been associated with a variety of skin disease conditions in several studies. Chronic dermal ER stress, caused by Darier’s disease and Ekpoma virus (EKV), triggers PERK-CHOP, which then activates the ER-mediated mitochondrial apoptosis pathway, allowing skin cells to die (46–50). Keratinization disorders and Keratosis linearis with Ichthyosis Congenita and Sclerosing Keratoderma syndrome (KLICK syndrome) are caused by ER stress that activates UPR signaling and promotes these disorders (51, 52). Patients with Vitiligo have a poor response to ER stress due to a deficiency in their melanocytes that exacerbates inflammation (53, 54). In this study, the effect of LSDV on ER stress-mediated apoptosis and autophagy was not adequately determined. Further research on ER stress-induced apoptosis and degradation may provide new insights into LSDV-infected cellular processes. Collectively, these findings reveal that LSDV infection triggers an ER imbalance, elicits ER ultrastructural abnormalities, and activates all three UPR sensors. Furthermore, LSDV infection is aided by PERK-activated translational repression and IRE1-mediated cleavage of XBP1, but ATF6 has a limited role in LSDV invasion. A possible therapy for LSDV infection is chemical suppression of particular UPR signals. These findings broaden our understanding of LSDV infection and provide potential LSDV molecular targets. The discovery of new insights into the molecular process of LSDV infection will aid in the development of antiviral medications.

## MATERIALS AND METHODS

### Cells, viruses, and animals

The primary bovine embryonic fibroblast (BEF) cells were a kind gift from Huaijie Jia. Madin-Darby bovine kidney (MDBK) cells were purchased from the China center for type culture collection (CCTCC) (Wuhan, China). The primary lamb testis (LT) cells stored in our lab were used to propagate and titrate the virus.

The virulent LSDV, LSDV/China/Xinjiang/2019, was provided by the Lanzhou Veterinary Research Institute, Chinese Academy of Agricultural Sciences. The strain was exposed and utilized in bio-safety level 3 (BSL-3) laboratories. After experiments, viral inactivation was performed by exposing LSDV in a 2% NaOH environment.

Six seronegative male Holstein cows aged six months were randomly divided into two groups (a Mock group and an LSDV-infected group) and separately housed in the BSL-3 lab. After a week of acclimatization, the cows were used in subsequent experiments.

### Antibodies

The primary antibodies involved in this research were GAPDH (ab181603, Abcam), IRE1 (ab37073, Abcam), phospho-IRE1 (ab124945, Abcam), XBP1 (ab220783, Abcam), ATF6 (ab122897, Abcam), GRP78 (3177S, CST), eIF2α (9722S, CST), phospho-eIF2α (3398S, CST), PERK (bs-2469R, Bioss), phospho-PERK (bs-3330R, Bioss), GRP78 (GB12098, Servicebio) and ORF122 (rabbit polyclonal antibody, kindly provided by Guohua Chen). The secondary antibodies were Goat Anti-Rabbit IgG H&L (HRP) (ab6721, Abcam), Goat Anti-Mouse IgG H&L (HRP) (ab205719, Abcam), Goat Anti-Mouse IgG Alexa Fluor® 488 (GB25301, Servicebio) and Goat Anti-Rabbit IgG Cy3 (GB21303, Servicebio).

### LSDV infection of Holstein cows *in vivo* and ethics statement

As described by the World Organisation for Animal Health (WOAH) (https://www.woah.org/en/disease/lumpy-skin-disease/) and in a previous study (55), cows in the LSDV-infected group were given multiple intradermal injections of a total of about 2-mL virus suspension. Likewise, cows in the MOCK group were administered intradermally with an equal volume of DMEM. Subsequently, the rectal temperature and characteristic phenotype (classic lumpy skin) were observed and recorded daily. On day 14 after administration, typical lumps were observed. Next, cows in both groups were sacrificed. Normal skin from the control, lumpy skin, and juxta-lumpy skin (5–10 cm from the lumps) from the LSDV-infected group were collected. Parts of samples were snap frozen in liquid nitrogen and stored at −80°C for making frozen sections and post-transcriptional protein assays, while others were removed and fixed in 4% paraformaldehyde (G1101, Servicebio) and 2.5% glutaraldehyde (G1102, Servicebio), respectively, for histological staining and transmission electron microscopy (TEM).

All animal experiments described above were performed in accordance with the instructions of Lanzhou Veterinary Research Institute (Chinese Academy of Agriculture Science) institutional animal care and the Good Animal Practice Requirements of the Animal Ethics Procedures and Guidelines of the People’s Republic of China.

### Hematoxylin & eosin (H&E) staining, immunohistochemistry (IHC), and indirect immunofluorescence (IFA)

Skin samples after fixation with paraformaldehyde underwent routine paraffin preparation. Briefly, transparency, paraffin immersion, embedding, and sectioning were conducted for obtaining 5-μm sections.

Parts of sections were stained with HE (G1005, Servicebio) according to the manufacturer’s instructions. Others sections were used for IHC staining. Briefly, after deparaffinization and rehydration, samples underwent antigen retrieval, peroxidase removal, permeabilization, blocking, antibody incubation, DAB development, and hematoxylin redyeing.

Skin samples frozen at −80°C were cut into 10-μm frozen sections. To evaluate the effect of LSDV infection on the GRP78 (Servicebio) and ORF122 distribution in the skin, samples were incubated with primary antibodies GRP78 and ORF122 from two different host sources. After that, secondary antibodies Alexa Fluor® 488 and Cy3 were labeled. Finally, DAPI (C0065, Solarbio) was used for nuclear labeling.

All sections mentioned above were scanned using a Pannoramic MIDI Digital Slide Scanner (3DHISTECH, Budapest, Hungary).

### Transmission electron microscope (TEM) observation

After fixing with 2.5% glutaraldehyde, skin samples were embedded in resin. Specimens were viewed with a HT7800 transmission electron microscope (Hitachi).

### Enzyme-linked immunosorbent assay (ELISA)

Serum samples from control cows and LSDV-infected cows were collected on day 0, day 3, day 6, and day 14 to identify LSDV infection. Subsequently, the competitive ELISA was performed according to the manufacturer’s instructions.

### Cell culture, virus infection, and chemical treatments

All cells were cultured in Dulbecco’s modified Eagle’s medium (DMEM) (Gibco) supplemented with 10% fetal bovine serum (FBS) (Gibco) and 1% penicillin streptomycin (Gibco) at 37°C and 5% CO_2_. When grown to approximately 70% confluence, MDBK and BEF cells were infected with LSDV at an MOI of 1 or mock infected with DMEM for 1 h. Subsequently, cells were washed with PBS and cultured in 2% FBS DMEM maintenance medium.

The three UPR inhibitors, GSK2606414 (a highly selective PERK inhibitor), 4μ8C (a highly potent IRE1 inhibitor), and Ceapin-A7 (a selective blocker of the ATF6α signal) were dissolved in DMSO. Both types of cells were cultured in 6-well plates (for WB) and 12-well plates (for IFA, virus titration, and extraction of viral genomes) and were pretreated with chemicals or DMSO for 24 h, followed by inoculation with LSDV as described above. Cells were harvested at 48 hours post-infection (hpi) for performing SDS-PAGE, western blotting, and cellular indirect immunofluorescence (IFA). Additionally, the supernatant was harvested for virus titration and extraction of viral genomes.

### Cell Viability Assay

Cell viability was detected using a Cell Counting Kit-8 (Beyotime, C0038, China) according to the manufacturer’s instructions.

### RT-qPCR

To confirm ER stress induction, MDBK and BEF cells were infected with LSDV (MOI=1) at different time points. Cells cultured in 6-well plates were collected at 12 hpi, 24 hpi, 48 hpi, and 96 hpi. Total cellular RNA was extracted with the TRIzol™ Reagent (15596018, Invitrogen) according to the manufacturer’s instructions. After that, total RNA underwent reverse transcription using a PrimeScript™ RT reagent Kit with gDNA Eraser (Perfect Real Time) (RR047A, Takara). The Bio-Rad iQ5 system was used to perform a qPCR procedure with the Hieff^®^ qPCR SYBR Green Master Mix (No Rox) (11201ES03, Yeasen). Relative mRNA transcriptional levels were normalized to GAPDH (an internal control). Primers involved in this research are listed in Table 1.

**TABLE 1.**
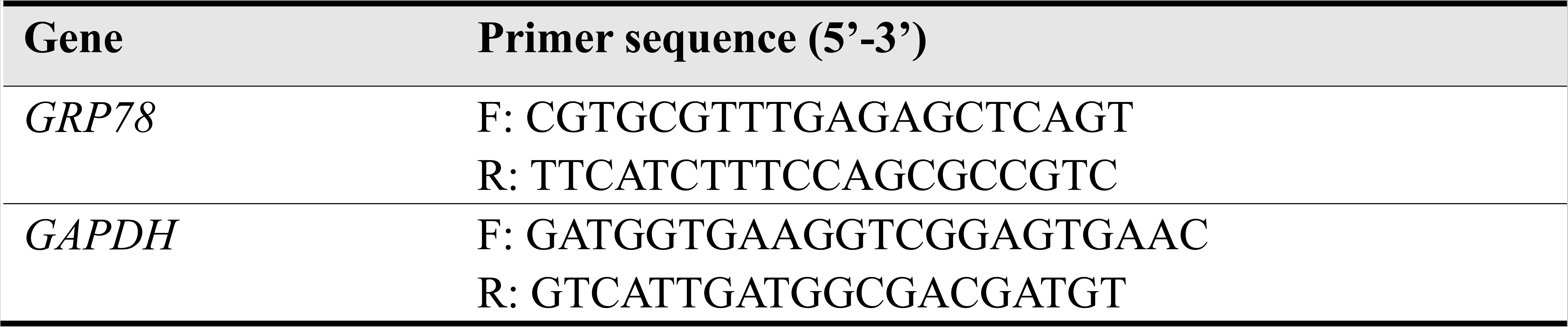
Primers used for qPCR analysis.

### Western blot

Tissue and cell samples were lysed by a RIPA lysis buffer (P0013B, Beyotime) supplemented with phenylmethylsulfonyl fluoride (PMSF) protease inhibitor (36978, Thermo) and phosphatase inhibitor cocktail A (50×) (P1082, Beyotime). Proteins with the same loading were separated by SDS-PAGE. Subsequently, the standard western blot procedure was performed. Briefly, proteins were sequentially subjected to PVDF membrane transfer, blocking, antibody incubation, and color development.

### Cellular indirect immunofluorescence (IFA)

Briefly, cells cultured in 12-well plates underwent fixation, permeabilization, blocking, primary antibody incubation, and Cy3 labeling. Images were collected using a DMI 6000B inverted fluorescence microscope (Leica).

### RNA interference

One control siRNA and nine UPR siRNAs were purchased from Genepharma (Shanghai, China). Their sequences of sense strands are presented in Table 2. The potency of siRNAs was assessed through western blots when cells were interfered for 72 h with different siRNAs. Next, MDBK and BEF cells cultured in 12-well plates were interfered for 24 h with the three representative UPR siRNAs, followed by inoculation with LSDV (MOI = 1) as described above. At 48 hpi, cells were harvested for performing cellular IFA. The supernatant was harvested for virus titration and extraction of viral genomes.

**TABLE 2.**
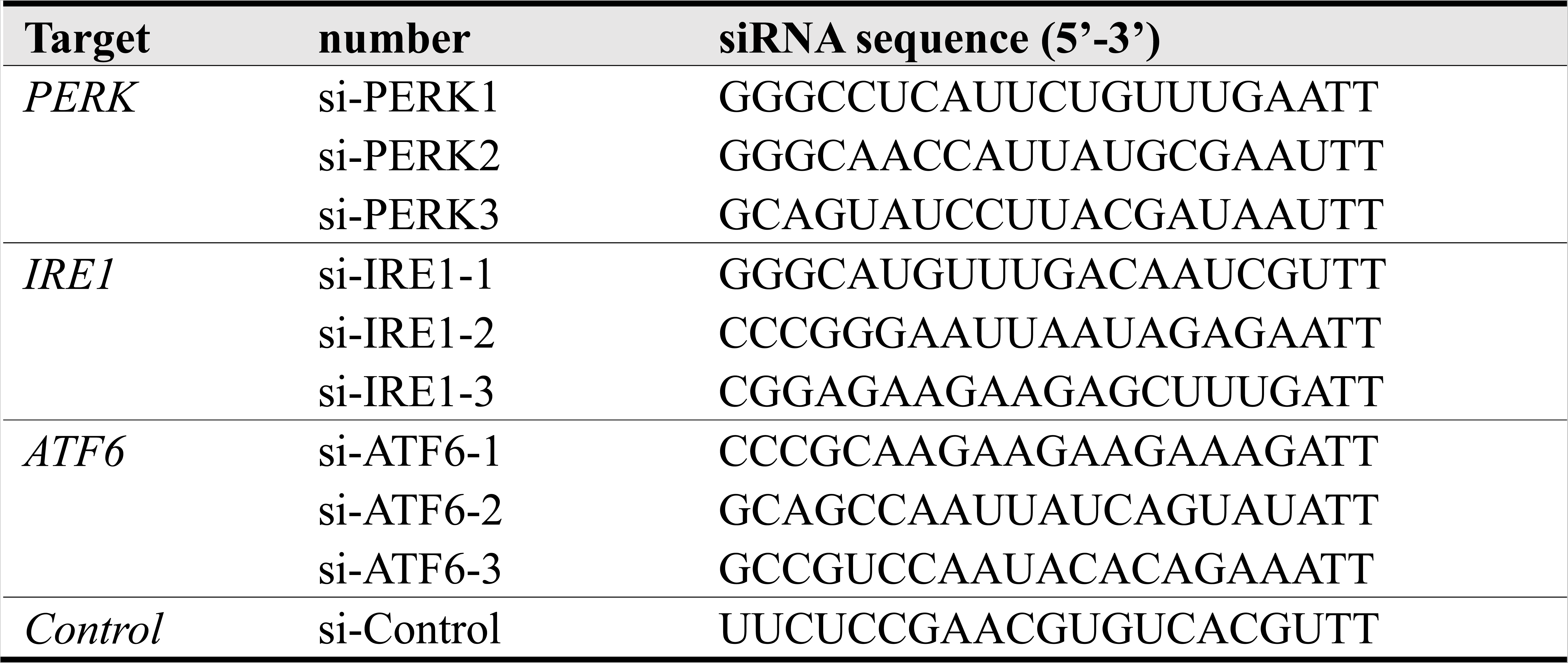
Sequences of siRNA sense strands used for UPR signal interference.

### Virus Replication Assay

After LSDV infection, siRNA interferon, or chemical pretreatments, MDBK and BEF cells were cultured in 12-well plates for two days. Subsequently, cells were subjected to three freeze-thaw cycle, and the cellular supernatant was collected. Virus genomes were extracted with an EasyPure^®^ Viral DNA/ RNA Kit (TransGen Biotech, Beijing, China) according to the manufacturer’s instructions. Subsequently, probe-based absolute quantitative PCR assays were performed as previously described (56). Copy numbers were examined to reveal the effects of various treatments on viral replication. Plasmid standards, probes, and primers were synthesized by Tsingke (Beijing, China). Prior to the experiment, the plasmid standard was identified (Fig. S3). Probe sequences and primer sequences are listed in Table 3.

**TABLE 3.**
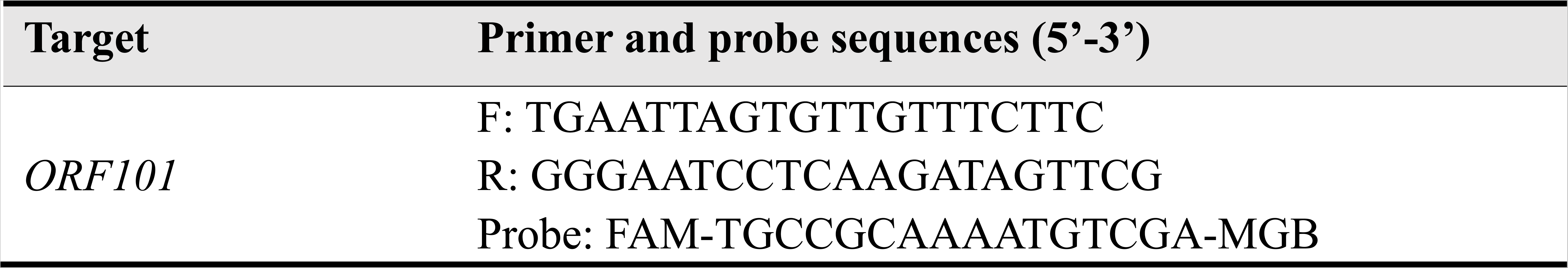
Primer and probe sequences used for virus absolute quantitation

### Virus titration

The supernatant was obtained as mentioned above. As per an OIE recommendation, the primary LT cells were used to titrate LSDV in 96-well plates. Briefly, serial 10-fold dilutions of the supernatant from 10^−1^ to 10^−10^ were added to 96-well plates growing 75% confluent LT cells. Cytopathic effects (CPE) were observed at 6 dpi to 10 dpi. The Kärber method was used to calculate virus titers expressed as TCID_50_/0.1 mL.

### Statistical analysis

Data were presented as mean and standard deviation (SD). One-way ANOVA and *t*-tests were performed for all statistical analyses using GraphPad Prism software. *P* values represent significant differences (*, *P* < 0.05; **, *P* < 0.01; ***, *P* < 0.001; ****, *P* < 0.0001), and ns indicates no significant difference.

## ACKNOWLEDGMENTS

This work was supported by the National Natural Science Funds for the National Natural Science Foundation of China (31972695). We thank LetPub (www.letpub.com) for its linguistic assistance during the preparation of this manuscript.

---

**Fig S1.**
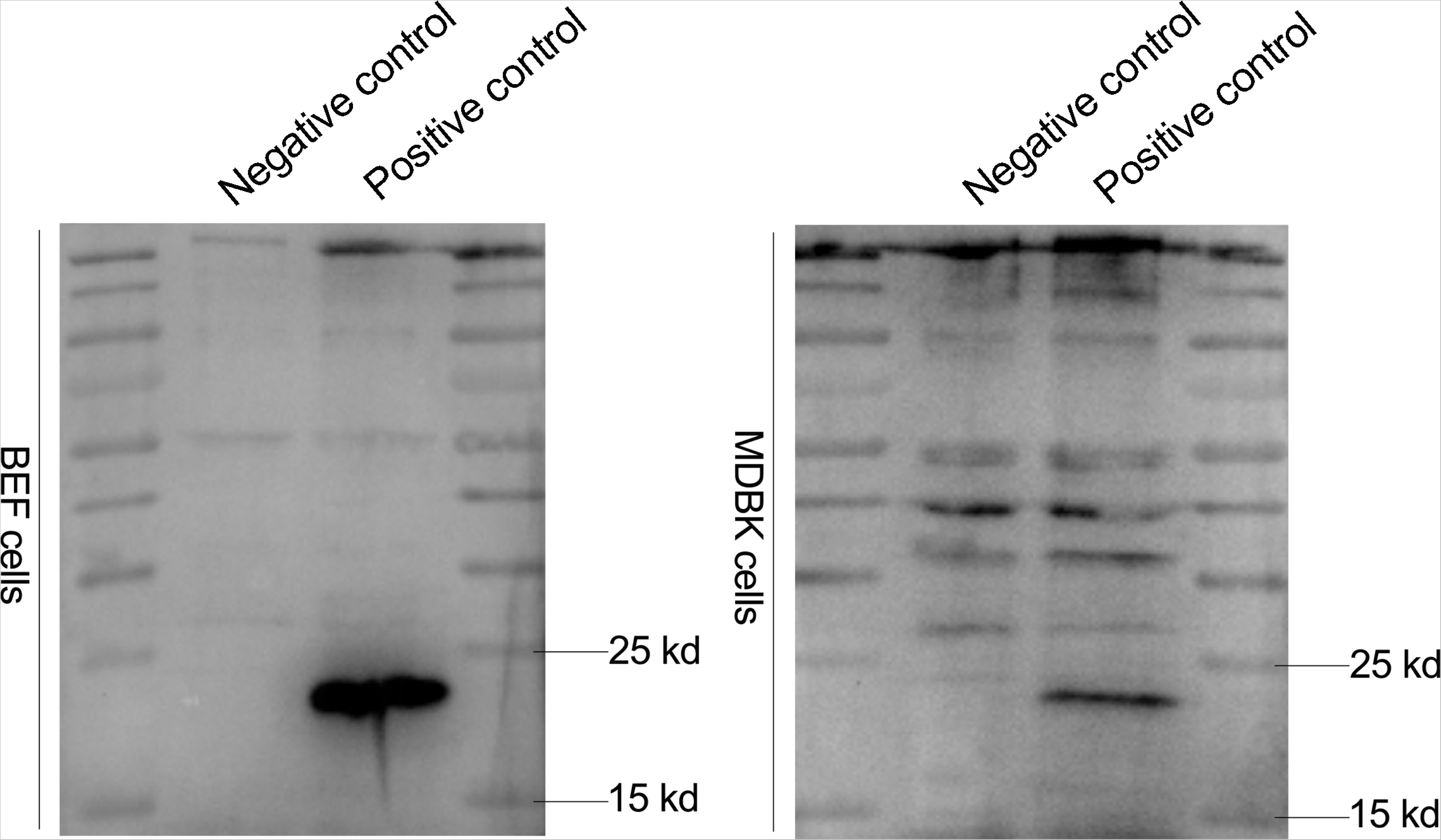

**Fig S2.**
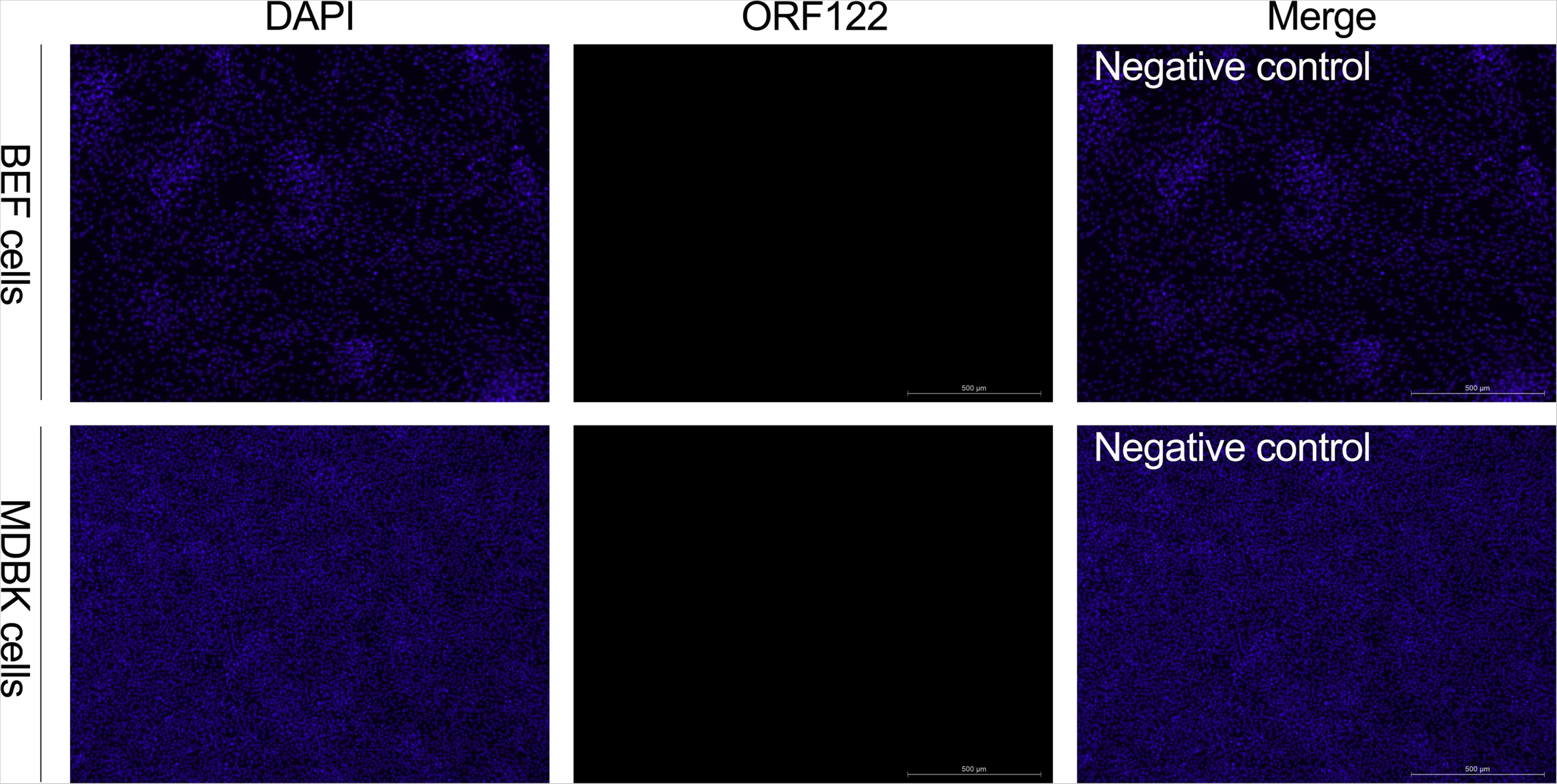

**Fig S3.**
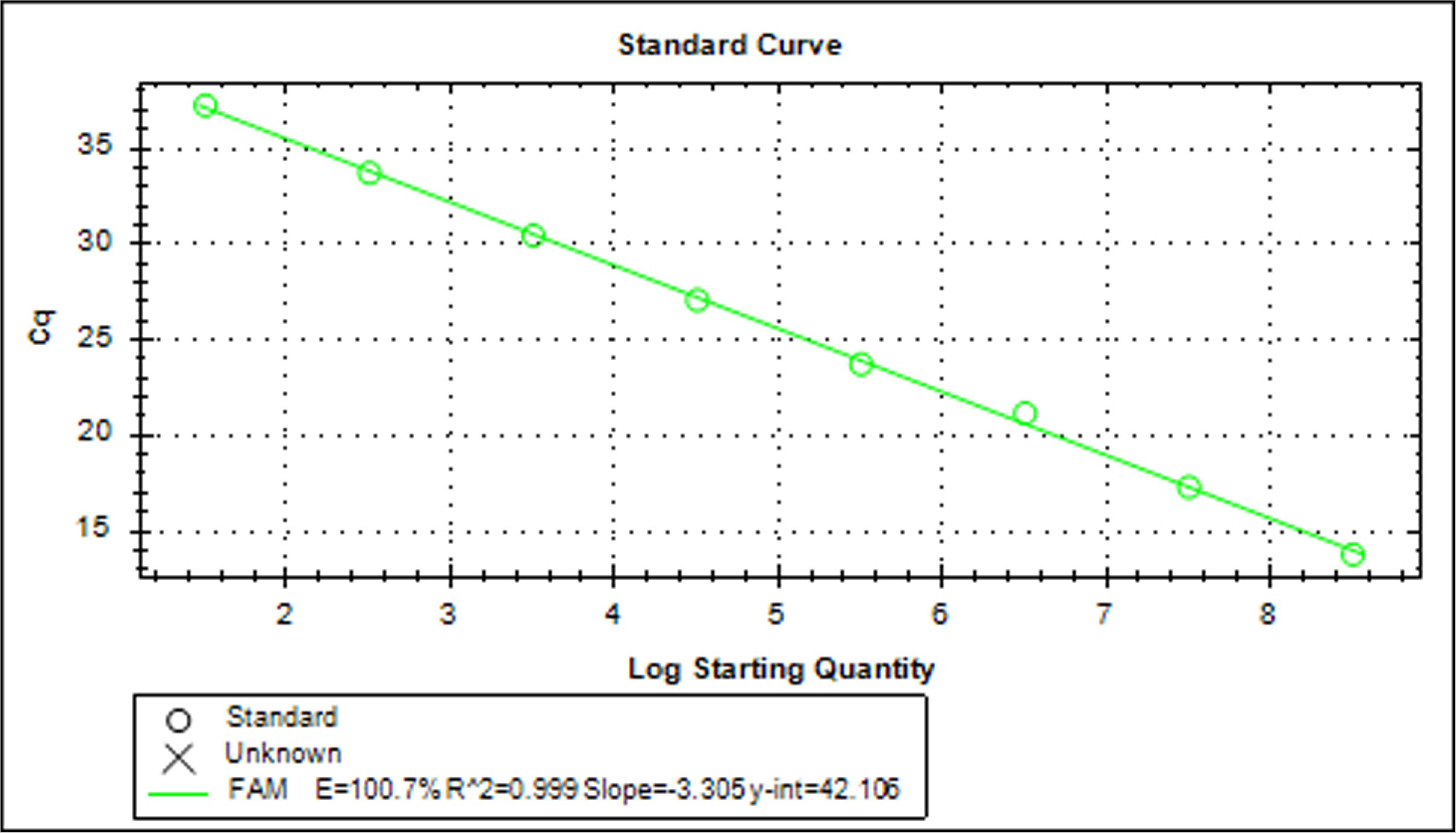

